# Stability of the Wheat Seed Mycobiome Across North Carolina’s Longitudinal Gradient

**DOI:** 10.1101/2024.02.22.581680

**Authors:** Lindsey E. Becker, Christine V. Hawkes, Ryan Heiniger, Marc A. Cubeta

## Abstract

Improving wheat yield and performance involves selecting varieties that are well adapted for a regional area. Although host genotype and environment are major factors that impact crop performance and resilience, less is known about the relative contribution and occurrence of wheat seed endophytic fungal communities across spatial and temporal scales. An increased understanding of composition and assembly of beneficial endophytic fungal communities across regional scales provides valuable insight into the stability of the endophytic seed mycobiome. Our aim in this study was to examine the relative contribution and impact of latitude and longitude gradients within North Carolina (NC) on wheat seed fungal community structure of two regionally adapted soft red winter wheat cultivars, Hilliard and USG 3640. We examined the endophytic wheat seed microbiome of the two winter wheat cultivars planted in official variety trials at five geographic locations across NC in 2021 and two geographic locations in 2022. ITS1 sequence-based analysis of surface disinfested wheat seeds was conducted to determine alpha and beta diversity. Species richness is influenced by geographical location, however wheat seed mycobiome community structure is stable across cultivars and years. Latitude and longitude contributed to the observed variation in wheat seed mycobiome structure, in addition to yield, seed moisture, and leaf nutrients. When surveying taxa present within all cultivars, geographical sites and years, *Alternaria* and *Epicoccum* spp. exhibited high relative abundance in the wheat seed mycobiome. Our results provide a comprehensive catalog of core fungal taxa well-adapted to diverse environments and conserved across wheat cultivars.

## INTRODUCTION

Recently, agricultural microbiomes, or microbial communities associated with crop plants, have been implicated in plant health, including disease resistance, drought tolerance, and flowering time (Adam et al. 2018; Bergna et al. 2018; Panke-Buisse et al. 2015; Benitez et al. 2017). Although root and leaf associated microbiomes have received much attention in cultivated crop species, seed communities merit further examination due to the potential for vertical transmission of microorganisms that are traditionally horizontally transmitted (Bergelson et al. 2019; Stopnisek and Shade 2021; Chaparro et al. 2014; Benitez et al. 2017; Johnston-Monje et al. 2016). Seeds represent a unique ecological niche within the plant and serve as a physical link between generations that could affect microbiome assembly if those taxa have priority in colonizing broader plant tissues upon germination. The seed microbiome has been shaped by domestication (Kim et al. 2020; Hassani et al. 2020), cultivar (Rybakova et al. 2017; Adam et al. 2018), seed origin (Wolfgang et al. 2020), disease (Rojas et al. 2020), geography (Klaedtke et al. 2016), among other factors. In wheat, the seed mycobiome is shaped by genotype (Latz et al. 2021) and influenced by pathogen infection (Bakker and McCormick 2019; Rojas et al. 2020). The development of the wheat seed over time substantially changes the composition of the seed mycobiome (Hertz et al. 2016). Understanding the assembly and sourcing of the seed microbiome as well as its contribution to the developing plant represents a key underpinning necessary to realize the benefits of the seed microbiome (Barret et al. 2015; Shade et al. 2017).

Plant associated fungi range in their interactions with host plants from commensal to pathogenic and mutualistic. Plants harbor distinct fungal communities (mycobiomes), with below and above ground tissues exhibiting differences in terms of composition and diversity (Gdanetz and Trail 2017; Latz et al. 2021). As a flowering plant undergoes reproduction, pathogenic and non-pathogenic fungi attempt to colonize the developing seed niche in waves of introductions from other plant organs and the surrounding environment (Hertz et al. 2016; Shade et al. 2017). Fungi present in the environment must bypass floral barriers, navigate beyond the seed coat, and avoid activating the immune system, by adapting to a rapidly changing environment as the seed matures (Nelson 2018; Vannette 2020). These barriers result in a small cohort of active seed fungal endophytes living within seed tissue (Newcombe et al. 2018). What is unclear is whether specific fungi are adapted to the seed environment and therefore always present or if the seed fungal community is shaped by fungal communities present in the local environment and host filtering.

To address these ideas, we tested whether the seed microbiomes were consistent across multiple sites and years, or instead tracked local conditions. We focused on wheat, an important C3 grass crop, because non-crop C3 grasses have developed fungal vertical transmission. We evaluated two wheat cultivars as previous studies suggested that the wheat seed microbiome might be genotype-specific (Latz et al., 2021). using a common garden experiment. Our experimental approach involved i); examining alpha diversity differences among sites and years, ii) examining whether beta diversity is structured by geography; iii) determining the relative influence of environmental factors on the seed fungal community, and iv) defining the core fungal taxa present across sites and years. By planting single-sourced seeds in five locations in North Carolina and over a period of two years, we provided high level detail of spatial and temporal influence on the wheat seed endophytic mycobiome.

## MATERIALS AND METHODS

### Plant Material

Wheat (*Triticum spp.*) has highly variable traits that translate into regional breeding success, with skilled wheat breeders selecting for many traits such as daylength, yield, milling quality, drought tolerance, and disease resistance (Borlaug 1983; Peng et al. 2011). Regionally-adapted cultivars are valued by growers as they are tailored for their specific growing conditions and needs. Cultivars are enrolled in field trials within and beyond their region to test cultivar performance in a variety of testing sites with known soil characteristics, climatic conditions, and disease pressure. These trials are conducted by land grant universities across the United States as Official Variety Trials (OVT). In North Carolina (NC), OVTs provide insight into cultivar performance for breeders and growers in different geographic regions across the state. Two winter wheat cultivars, Hilliard and USG 3640, were selected for this study. Hilliard is an awned soft red winter wheat, heading mid-season, widely adapted for the eastern US and released by VA Tech in 2015. Hilliard is desirable due to its consistent high grain yield and moderate resistance to leaf and stripe rust, Fusarium head blight, and leaf and glume blotch diseases (Griffey et al. 2020). USG 3640 is a soft red winter wheat, heads mid-season and is awned, released by University of Georgia soft red winter wheat breeding program and licensed to Uni South Genetics as USG 3640. USG 3640 is a high yielding cultivar well-suited to the southeastern US with moderate resistance to powdery mildew, leaf and stripe rusts (Mergoum et al. 2022). Hilliard has performed well in North Carolina State University Official Variety Trials (OVT) (Above-Average-Multiple-Year_Commercial-2020_22.pdf n.d.). Untreated USG 3640 and Hilliard seeds were sourced from a single field at the Midpines USDA-ARS Research Station in Raleigh, NC in 2020 and 2021.

### Field Sites

Sites for this study are associated with the OVT conducted in 2021 and 2022. Each field location represents a unique wheat-growing region in North Carolina, spanning various soil types that include: Rains sandy loam, Nahunta very fine sandy loam, Lloyd clay loam, Badin channery silt loam, and Portsmouth fine sandy loam (Table S1). The sites are located in major geographic wheat production regions within NC; the Coastal Plains, Tidewater, and Piedmont. The sites vary slightly in their fertilizer needs and soil characteristics (Soil-and-Cultural-Practice-Table-2021.pdf n.d.; Soil-and-Cultural-Practice-Table-2022.pdf n.d.). Site location, soil reports and further details are listed in Table S1. Lenoir, Robeson, Rowan, Union, and Washington county sites were sampled in 2021 (year 1). Rowan and Union were additionally sampled in 2022 (year 2). The five sites range geographically from 34.522329 to 35.849167 ° latitude and −80.624 to −76.656 ° longitude (Table S1). Locations of each site are illustrated in Fig. 1.

**Figure 1.**
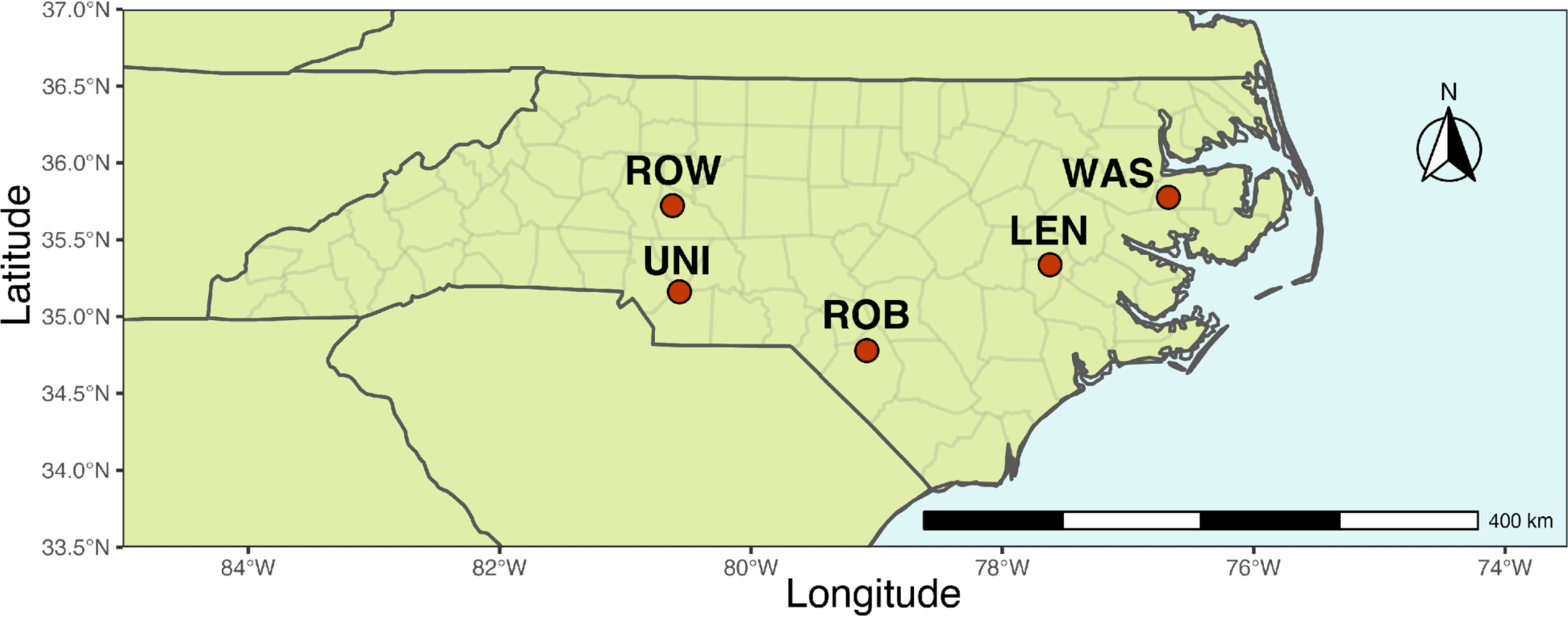
Map of North Carolina State University Official Variety Trial locations within North Carolina. ROW=Rowan, UNI= Union, ROB=Robeson, LEN=Lenoir, WAS=Washington.

### Experimental Design

Each site was planted with five replicate plots of each cultivar arranged in a complete randomized block design. Individual plots contained seven rows, with 7.5 in of space between each row and were 25 ft long. Each plot was planted at a rate of 26 seeds per row-foot or 1.8 million seeds per acre. Cultural practices, such as field plot preparation, date of planting, and fertilization were performed according to standard farming practices and were uniform for both cultivars at all OVT sites. Prior to planting each site, soil samples were obtained from the plots and fertilizer and lime applications were made accordingly. Wheat trials were designed to evaluate cultivar performance in the absence of fungicide sprays and seed treatment.

### Sampling

At harvest, each plot was individually harvested with a combine, bagged and analyzed for weight (kg), yield (estimated bushels/acre), test weight (kgs/bushel), and moisture (%). We consistently selected three plots per site for each cultivar. A total of 50 g of seeds from each selected plot were stored at 4 ℃ for further analysis.

### Seed processing

To examine the endophytic seed mycobiome, twelve seeds from each 50g seed selection per plot, field site, cultivar, and year were surface disinfested by immersing seeds for 30s in 95% EtOH, 2 min in 4% NaOH with 0.02% Tween 20, 30 s in 95% EtOH, 30s in sterile H_2_O, and 5s in sterile H_2_O (Verma et al. 2019). To test the success of seed surface disinfestation, we rolled a subset of disinfested seeds on malt extract nutrient agar by shaking the plate for 30 s and subsequently removing seeds (BD Difco, Franklin Lakes, NJ). Plates were incubated at room temperature (25 ℃). Plates were assessed for visible microbial growth over a 7-day period and no growth was observed. Separately, four disinfested seeds per plot free of visible damage or infestation were selected for individual seed DNA extractions. Each seed was aseptically placed in an individual bead-beating tube (Macherey Nagel, Düeren, Germany) and shaken for 30 s with a Bead Ruptor to homogenize the sample (Omni, Kennesaw, GA).

### DNA extraction and Library preparation

Genomic DNA was extracted from each homogenized sample using the Plant II Nucleospin kit (Macherey Nagel, Düeren, Germany). The DNA extraction method was adapted for fungal DNA by including an additional chloroform step prior to filtration per manufacturer’s suggestion. Extracted DNA for each sample was quantified using Qubit Fluorometer (Invitrogen, Carlsbad, CA) and stored at −20 ℃. To examine the mycobiome composition with metabarcoding, we utilized benchmarked and wheat leaf validated Internal Transcribed Spacer (ITS) 1 region primers (ITS-1F_KYO2: TAGAGGAAGTAAAAGTCGTAA and ITS86R: TTCAAAGATTCGATGATTCAC) (Scibetta et al. 2018). We optimized the recommended polymerase chain reactions (PCR) cycling conditions and performed a touchdown PCR in a T100 thermocycler (Bio-Rad, Hercules, CA). The total reaction volume was 50 µl which contained 2 µl of genomic DNA at 50 ± 25 ng/µl, 25 µl DreamTaq PCR Master Mix 2X (Thermo Fisher Scientific, Waltham, MA), 2.5 µl of each primer (10 µM), and 18 µl of sterile H_2_O. PCR cycling conditions included two phases, an initial touchdown phase which starts with denaturation at 95 °C for 3 min; followed by 15 cycles of denaturation at 95 ℃ for 30 s, annealing at 63 ℃ for 45 s, and extension at 72 ℃ for 1 min, with the annealing temperature decreasing by 1 ℃ for all 15 touchdown cycles. The second phase consisted of 20 cycles of denaturation at 95 ℃ for 30 s, annealing at 56 ℃ for 45 s, extension at 72 ℃ for 1 min, with a final extension at 72 ℃ for 5 min (Korbie and Mattick 2008). PCR products were examined for successful amplification by electrophoresis in 2% agarose gels and visualized with ethidium bromide staining. A 100 bp DNA ladder was used to determine amplicon size (Scibetta et al. 2018). Library preparation and sequencing were performed at the Michigan State Genomics Core facility. PCR products were batch normalized with SequalPrep DNA Normalization plates (Invitrogen, Carlsbad, CA) and products were pooled and a round of quality control was performed prior to loading each batch into two separate Illumina MiSeq version2 Standard flow cells. Sequencing was performed in 2 x 250 bp paired-end format (Illumina, San Diego, CA). A total of 168 samples were sequenced on two separate sequencing runs at a similar read depth. The sequence data for both runs were combined prior to downstream analysis. Raw sequencing data was accessioned in the NCBI SRA database under bioproject PRJNA1077707.

### Bioinformatics

To analyze raw sequence reads, we utilized DADA2 ITS Pipeline Workflow 1.8 with the following parameters for filtering: maxN=0, maxEE=c(2,2), truncQ=2, minLen=50, rm.phix=TRUE and truncLen=c(190,150). Sequence tables were merged, prior to error rates and chimeric reads filtering as standard in DADA2 ITS Workflow (Callahan et al. 2016). Lulu curation was performed to remove spurious Amplicon Sequence Variants (ASVs) from the dataset. Spurious reads were incorporated into more abundant ASVs to prevent inflation of diversity estimates (Frøslev et al. 2017). Taxonomy assignment for curated ASVs was performed in DADA2 using the UNITE ITS database Version 9.0 (Abarenkov et al. 2010). A total of 167 samples passed the quality filtering parameters and were used for downstream analysis. Due to the low complexity of the wheat seed mycobiome and prior Lulu curation, ASVs with less than 5 reads in our dataset were removed for beta diversity estimates.

### Statistical analysis

The R environment (version 4.2.2) was used for statistical analysis and data visualization (R Core Team 2018). Plots were produced using ggplot2 (version 3.4.0), phyloseq (version 1.42.0), phylosmith (version 1.0.6), and vegan (version 2.6-4) (Wickham 2016; McMurdie and Holmes 2013; Smith 2019; Oksanen et al. 2022). The OVT site map was generated with the ggplot2 and sf packages (version 1.0-9) (Pebesma 2018). Alpha diversity was calculated with phyloseq for Shannon diversity and observed richness. An analysis of variance (ANOVA) was performed on Shannon diversity estimates and Tukey HSD groupings were assigned to relevant variables using vegan. For beta diversity analyses, the centered log-ratio (CLR) transformation was performed on the ASV abundance table using vegan. The Euclidean distance matrix was calculated with the CLR transformed abundance estimates. PERMANOVA analysis was conducted on CLR-transformed abundance and the Euclidean distance matrix to examine differences in fungal communities using RRPP. Fungal community structure was visualized using principal co-ordinate analysis (PCoA) ordination plots. Vegan was employed to visualize fungal community structure and association with environmental and physiological variables in a post-hoc redundancy analysis (RDA). Phylosmith was utilized to construct a heat map for 30 most abundant ASVs across all sites and years. Vegan and ggplot2 were used to compute and plot abundance and occupancy plots (Shade and Stopnisek 2019). The rare_curve function in the vegan package was deployed to calculate rarefaction curves for all samples and visualize sampling depth.

## RESULTS

### Amplicon sequence variants and alpha diversity

A total of 168 samples of wheat seeds yielded 8,745,028 ITS1 reads initially, with an average of 103,491 reads per sample. Using the DADA2 ITS Workflow to filter out low quality reads, 7,058,341 reads remained. After denoising, forward and reverse reads were merged for a total of 6,432,827 reads, followed by removal of chimeric reads to yield a total of 6,084,388 reads with an average 32,216 reads/sample. Three hundred and ninety fungal Amplicon Sequence Variants (ASVs) were included in the complete dataset (Callahan et al. 2016). To evaluate sequencing depth, we examined rarefaction curves, confirming that all individual seed samples reached plateau with a minimum read depth of 8,076 reads (Fig. S1).

Shannon diversity estimates of alpha diversity ranged from 0.024 to 1.534 across all samples and observed species richness ranged from 3 to 19 ASVs per seed, with an average of 9 ASVs per seed. At the cultivar level little variation was observed between Hilliard and USG 3640 in years 1 and 2 (Fig. 2A and 2C). A significant difference in alpha diversity among sites (P = 0.004) was observed in year 1(Fig. 2B). Seeds sampled from wheat plants grown at the Washington field site exhibited the highest alpha diversity compared to the Robeson and Rowan locations, which grouped together based on Tukey HSD post-hoc analysis. No significant differences in alpha diversity was observed among cultivars or by location across years for Rowan and Union sites (Fig. 2D).

**Figure 2.**
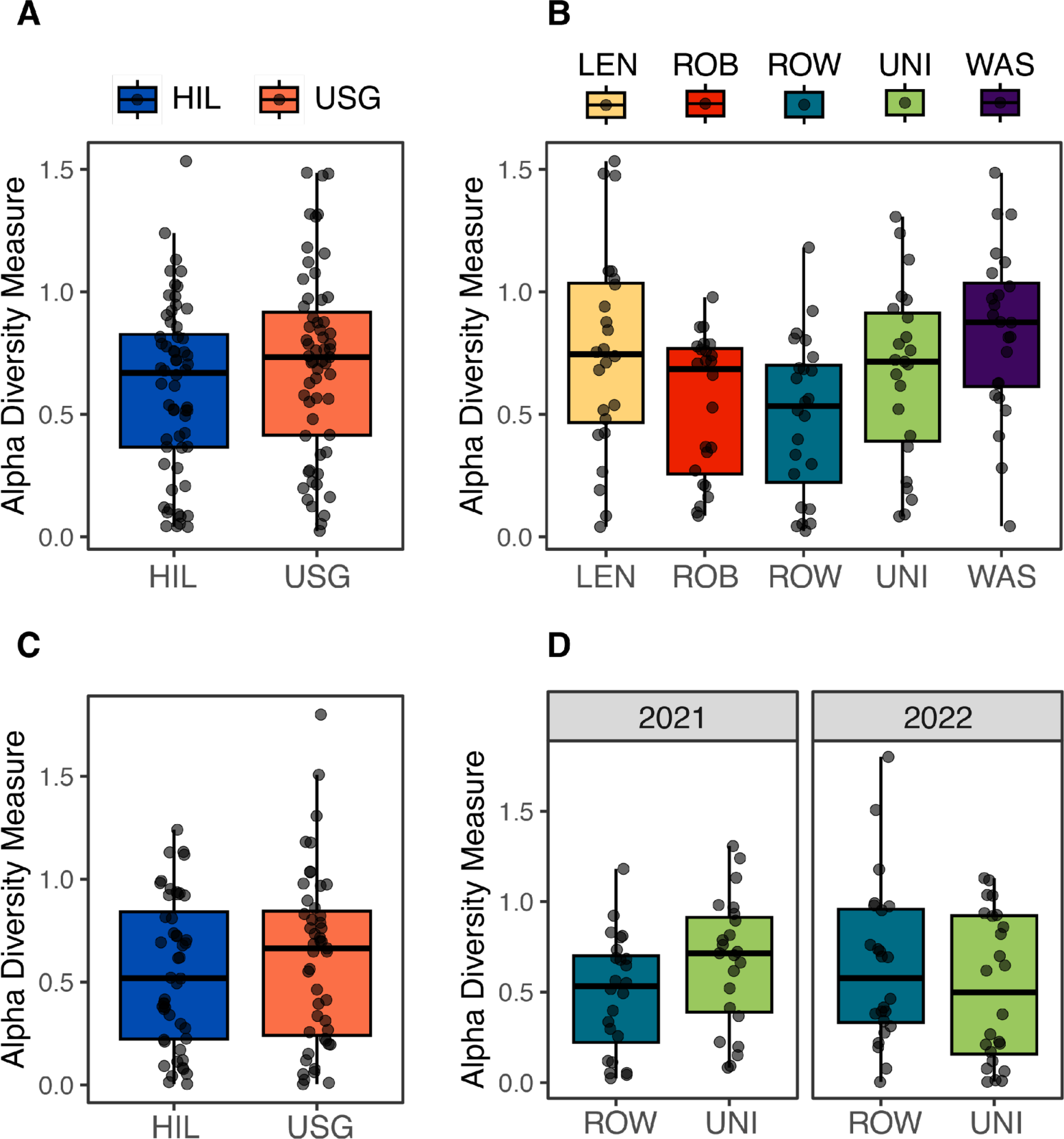
Alpha diversity of fungi associated with wheat seeds in Official Variety Trials estimated by Shannon Diversity. Comparisons of Shannon Diversity carried out using analysis of variance (ANOVA) by A) Cultivar for 2021 (year 1), B) Sites for year 1, C) Cultivar for 2022 (year 2), and D) Rowan and Union sites across years 1 and 2 (*P* < 0.05).

### Fungal community composition

To investigate the fungal community composition of our wheat seed samples, the relative abundance of the 20 most abundant taxa associated with year 1 (Fig. 3A) and Rowan and Union sites in 2021 and 2022 (Fig. 3B) was determined. The most abundant ASVs sampled from wheat seeds, belonged to two classes (Dothiedeomycetes and Sordariomycetes) within the Ascomycota, with *Alternaria* and *Epicoccum* representing the most abundant taxa sampled in 2021 (Fig. 3A) and for 2021 and 2022 at the Rowan and Union field sites (Fig. 3B).

**Figure 3.**
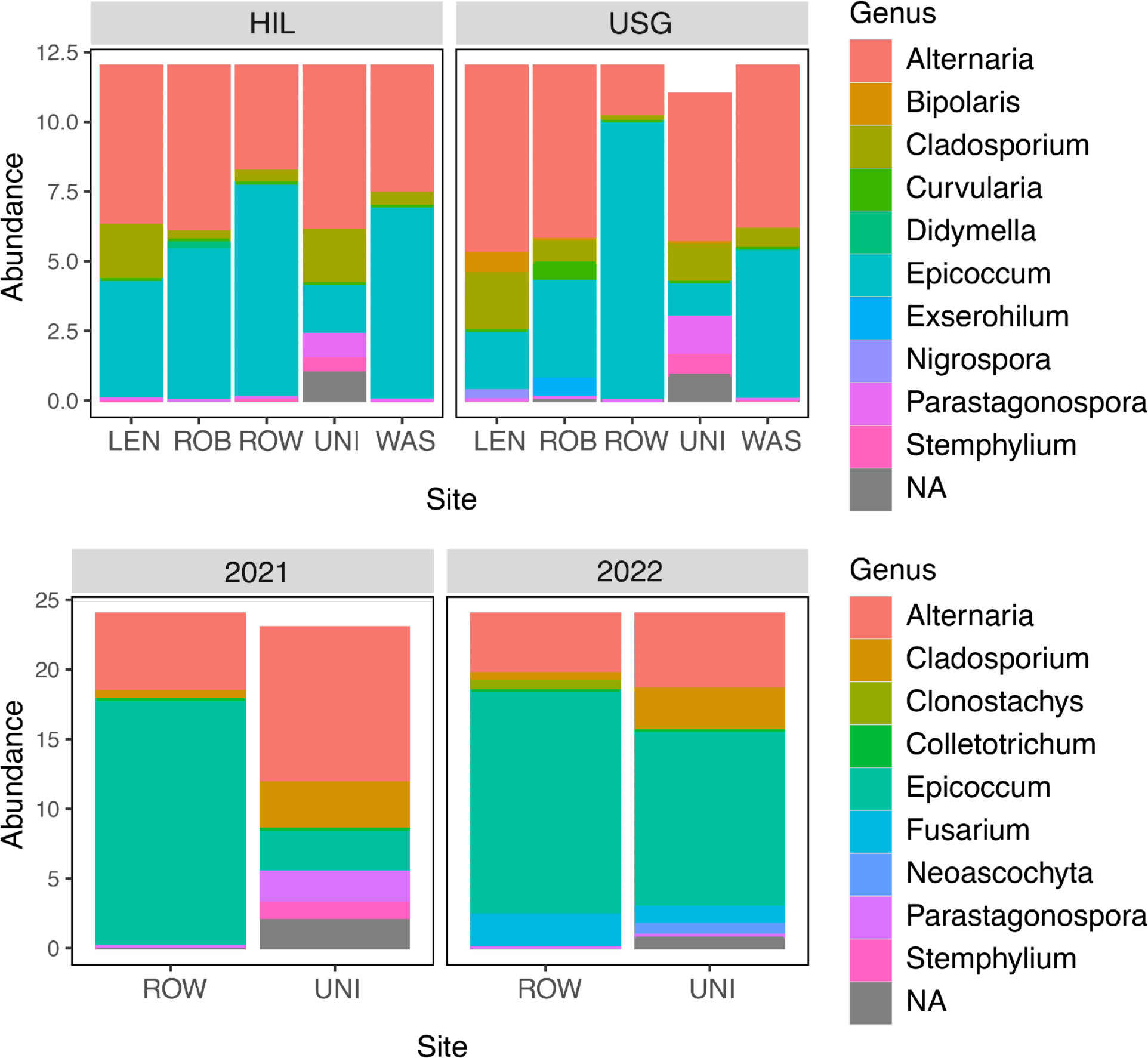
Relative abundance of fungal genera for fungal metabarcoding analysis A) Relative abundance of fungal genera by cultivar and site in year one. B) Relative abundance of fungal genera in Rowan and Union sites across years 1 (2021) and 2 (2022).

In Year 1, no variation in fungal community composition of seeds among sites (PERMANOVA, df= 4, *F value* = 0.78, *R^2^ =* 0.03, *P* value = 0.82) (Fig. 4A) or cultivars (PERMANOVA, df= 4, *F value* = 0.53, *R^2^ =* 0.004, *P* value = 0.85) was detected (Fig. 4A). Significant variation (PERMANOVA, df= 1, *F value* = 2.21, *R^2^ =* 0.025, *P* value = 0.006) was observed for fungal community composition among the Rowan and Union sites across years. However there was no variation observed by year (PERMANOVA, df= 1, *F value* = 0.94, *R^2^ =* 0.01, *P* value = 0.3) or cultivar (PERMANOVA, df= 1, *F value* = 1.21, *R^2^ =* 0.01, *P* value = 0.26) (Fig. 4B). Seed mycobiome variation at the plot level within each field site was further investigated for 2021 across all sites (Fig. S2A) and for Rowan and Union sites by each year (Fig. S2B). Each plot contained large amounts of variation, but no plot discernably separated from other plots based on PCoA analysis. These results suggest that single seed variation of individual samples likely contributes to the observed variation at the plot and site level, which may partially explain why differences in the fungal community were not observed among sites.

**Figure 4.**
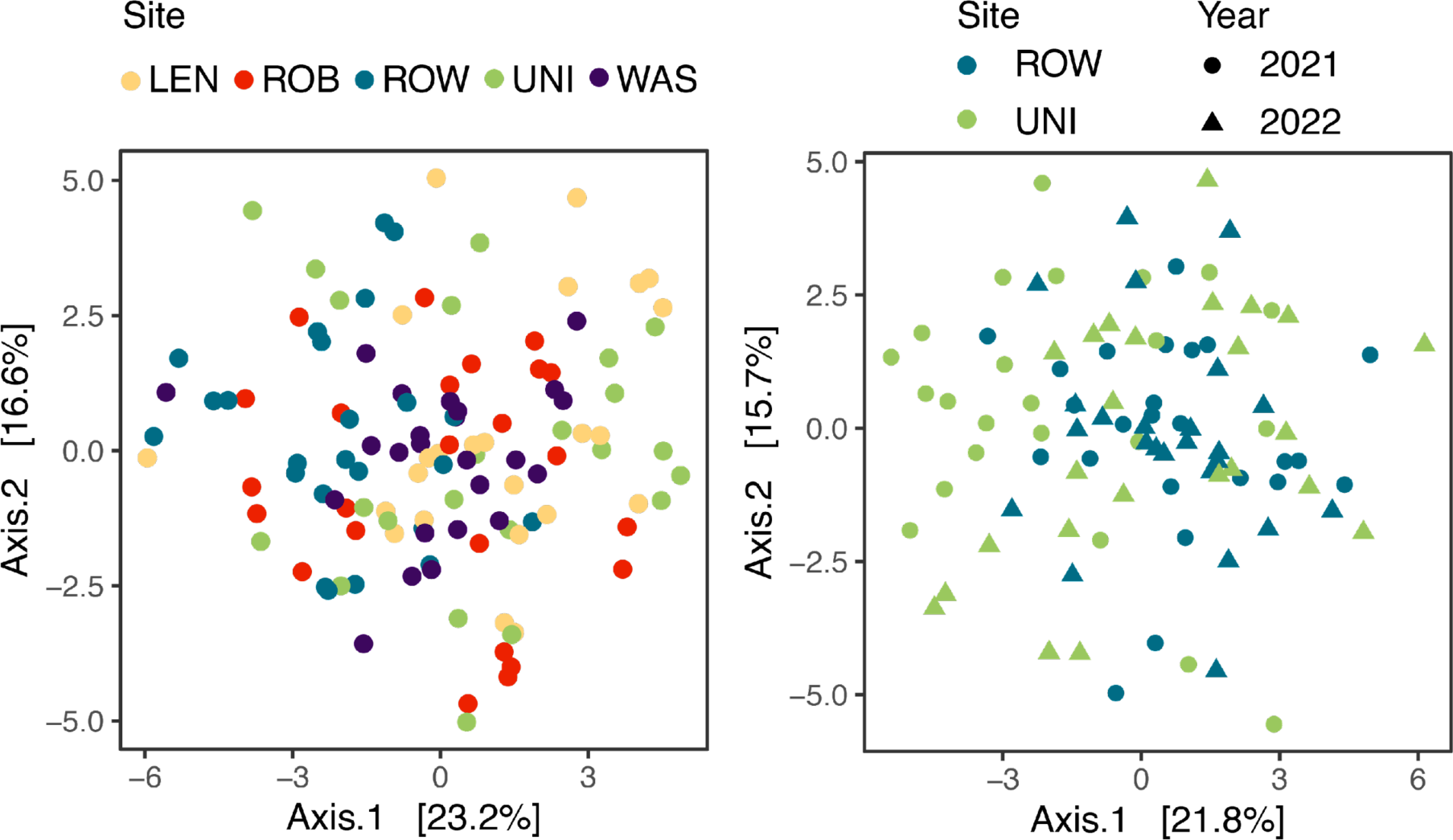
Principal Co-ordinates Analysis (PCoA) plot based on Euclidean Distances to examine structure of the wheat seed mycobiome. A) Seed mycobiome structure by site for year 1 (2021). B) Seed mycobiome structure for Rowan and Union sites over years 1 (2021) and 2 (2022). The first two components (PC1 and PC2) in 2021 accounted for 23.2 and 16.6% of total variation. PCoA analysis of Rowan and Union site ASV data indicated that PC1 accounted for 21.8% and PC2 contributed to 15.7% of variation observed.

To investigate the fungal community composition in the context of environmental variables, redundancy analysis (RDA) was used to visualize the relative strength of environmental factors and individual ASVs on community composition. Based on RDA, environmental factors accounted for 14.08% of variance for 2021 at five sites and 10.15% for Rowan and Union sites in 2021 and 2022. Environmental and physiological variables used for RDA included yield, latitude, longitude, seed moisture, rainfall, and leaf nutrients. In 2021, post-hoc analysis on Euclidean-distance PCA revealed that latitude (*R^2^ =* 0.09, *P* value = 0.02), seed weight (*R^2^ =* 0.08, *P* value = 0.03), yield (*R^2^ =* 0.09, *P* value = 0.03), leaf N (*R^2^ =* 0.08, *P* value = 0.01), leaf Mn (*R^2^ =* 0.06, *P* value = 0.04), and leaf K (*R^2^ =* 0.17, *P* value = 0.01) contributed to the seed mycobiome composition (Fig. 5A). Additionally ASVs 1, 2, 4, 5, and 6 were all noted to drive variation based on RDA. For example, ASV 1 (*Alternaria alternata*) was associated with longitude, ASV2 (*Epicoccum nigrum*) with leaf sulfur and iron content, ASV 4 (*Cladosporium* sp.) with seed moisture, ASV 5 (*Cladosporium herbarum*) and 6 (*Parastagonospora nodorum*) with leaf calcium content (Fig. 5A). For seed sampled from plots at the Rowan and Union sites across both years, post-hoc analysis of PCA revealed that latitude (*R^2^ =* 0.15, *P* value = 0.01), longitude (*R^2^ =* 0.15, *P* value = 0.01), and seed moisture (*R^2^ =* 0.10, *P* value = 0.02) were significant variables in shaping the seed mycobiome composition for the Rowan and Union sites (Fig. 5B). Within the Rowan and Union sites, ASVs 1, 2, 3, 5, 6, and 14 were noted as driving variation in the seed mycobiome. ASV 1 (*Alternaria alternata*), ASV 5 (*Cladosporium herbarum*), and ASV 6 (*Parastagonospora nodorum*) were associated with test weight and rainfall, while ASV 2 (*Epicoccum nigrum*) with latitude, ASV 3 (*Epicoccum nigrum*) with seed weight, and ASV 14 (*Fusarium chlamydosporum*) associated with seed moisture (Fig. 5B).

**Figure 5.**
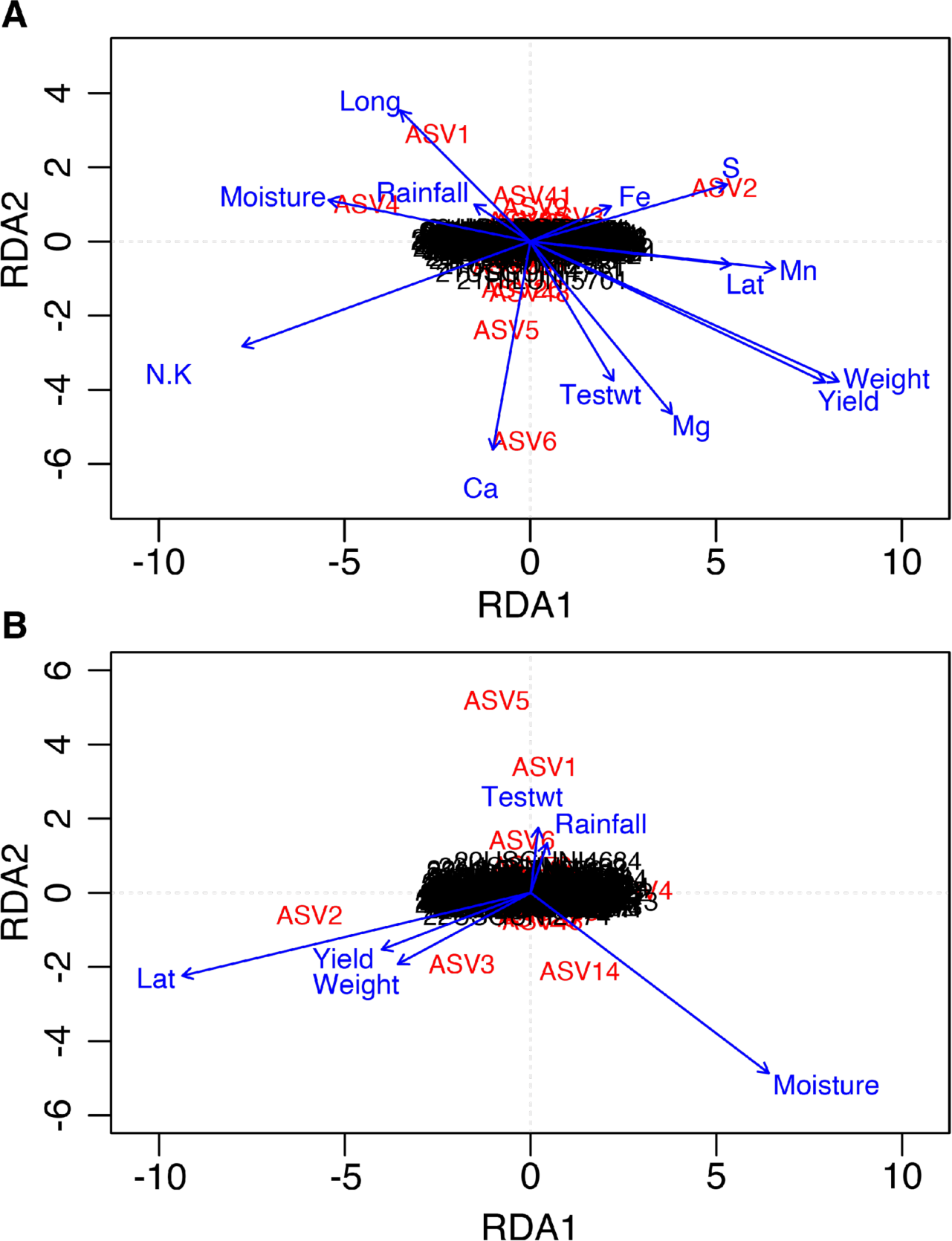
Principal Component Analysis (PCA) plot with Redundancy Analysis for environmental variables. A) Year 1 (2021) redundancy analysis with variables including latitude, longitude, yield, and leaf nutrients. B) Rowan and Union redundancy analysis for years 1 (2021) and 2 (2022) with variables including latitude, longitude, and yield.

To visualize the relative abundance of the top 30 ASVs for all 2021 field sites and the Rowan and Union field sites in 2021 and 2022 we utilized heatmaps (Figs. 6A and 6B). *Alternaria*, *Epicoccum*, and *Cladosporium* represented the most abundant genera across sites and within sites for 2021 and Rowan and Union sites in 2021 and 2022. Ascomycota genera were the most abundant, while three Basidiomycota genera (*Bullera*, *Filobasidium*, and *Papiliotrema*) were also present in both data sets. Site level variation was observed for less abundant genera *Parastagonospora*, *Stemphylium*, and *Filobasium*, while within site variation was observed for *Curvularia*, *Fusarium*, and *Pyrenophora* in 2021. An examination of Rowan and Union sites in 2021 and 2022 revealed that *Parastagonospora*, *Clonostachys* and *Neoascochyta* were variable between and within sites. The heatmap highlights the importance of comprehensive sampling at both plot and site levels as rare fungal taxa present in the seed may be plot and/or sample-specific. To examine the mean relative abundance and occupancy, four of the most abundant ASVs that occupied more than 75% of the seed samples across all field sites in 2021 and the Rowan and Union sites in 2021 and 2022 were *Alternaria alternata* (ASV 1), *Epicoccum nigrum* (ASV 2), and *Cladosporium* spp. (ASVs 4 and 5) (Figs. 7A and 7B). Genera at occupancy levels between a 25 and 75% of samples included ASV 3, 6, 33, 48, and 55 for 2021, representing *Epicoccum nigrum* (ASV 3), *Parastagonospora nodorum* (ASV 6), *Filobasidium* sp. (ASV 33), *Bullera* sp. (ASV 48), and *Papiliotrema* sp. (ASV 55).

**Figure 6.**
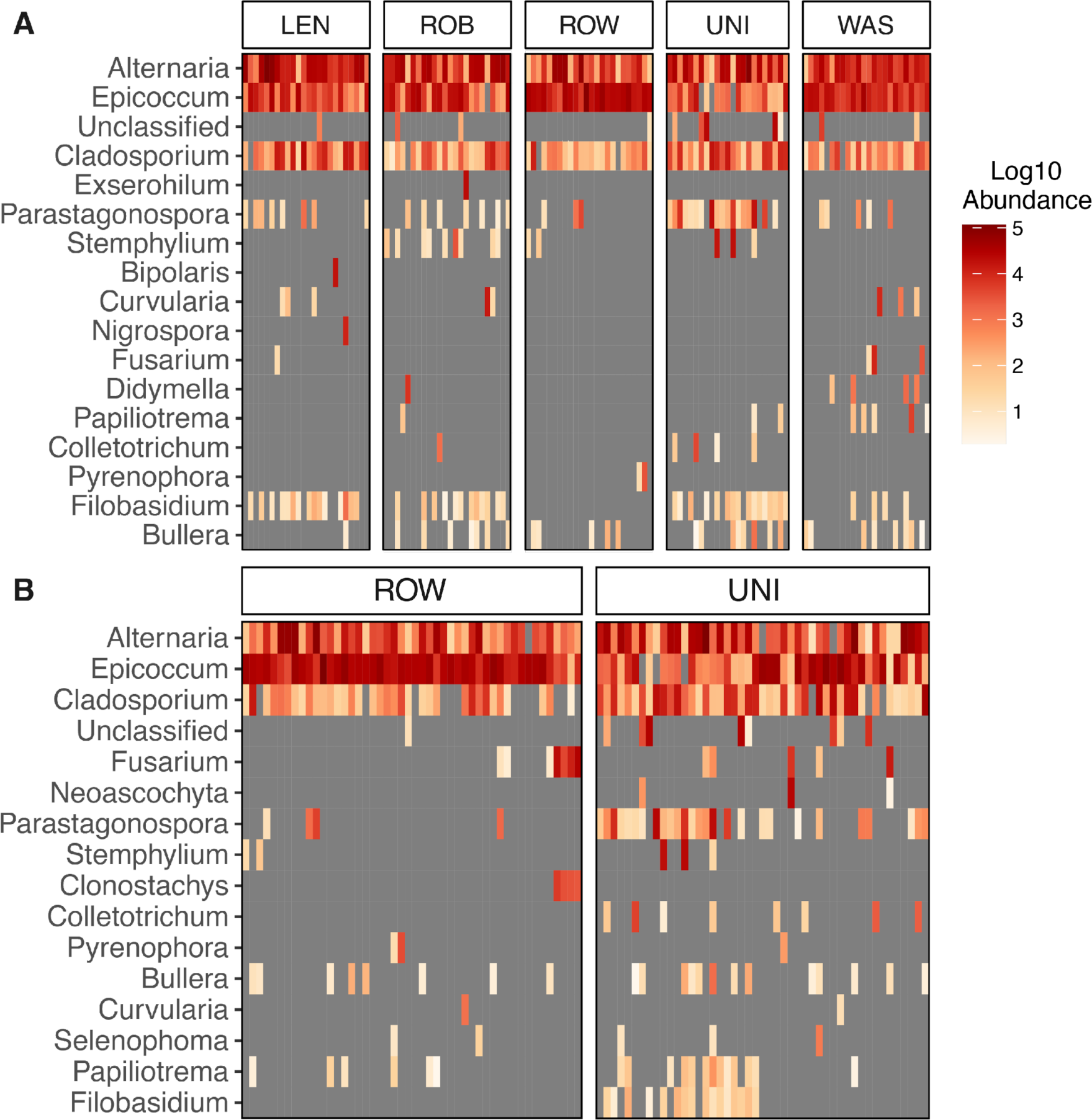
Heatmap of 30 most abundant ASVs classified at the genus level for A) sites surveyed in year 1 (2021) and B) Rowan and Union sites across two years (2021-2022). Grey color indicates absence of taxa within an individual sample. White to red color gradient indicates Log_10_ abundance of genera within a sample. Each vertical line represents an individual seed sample.

**Figure 7.**
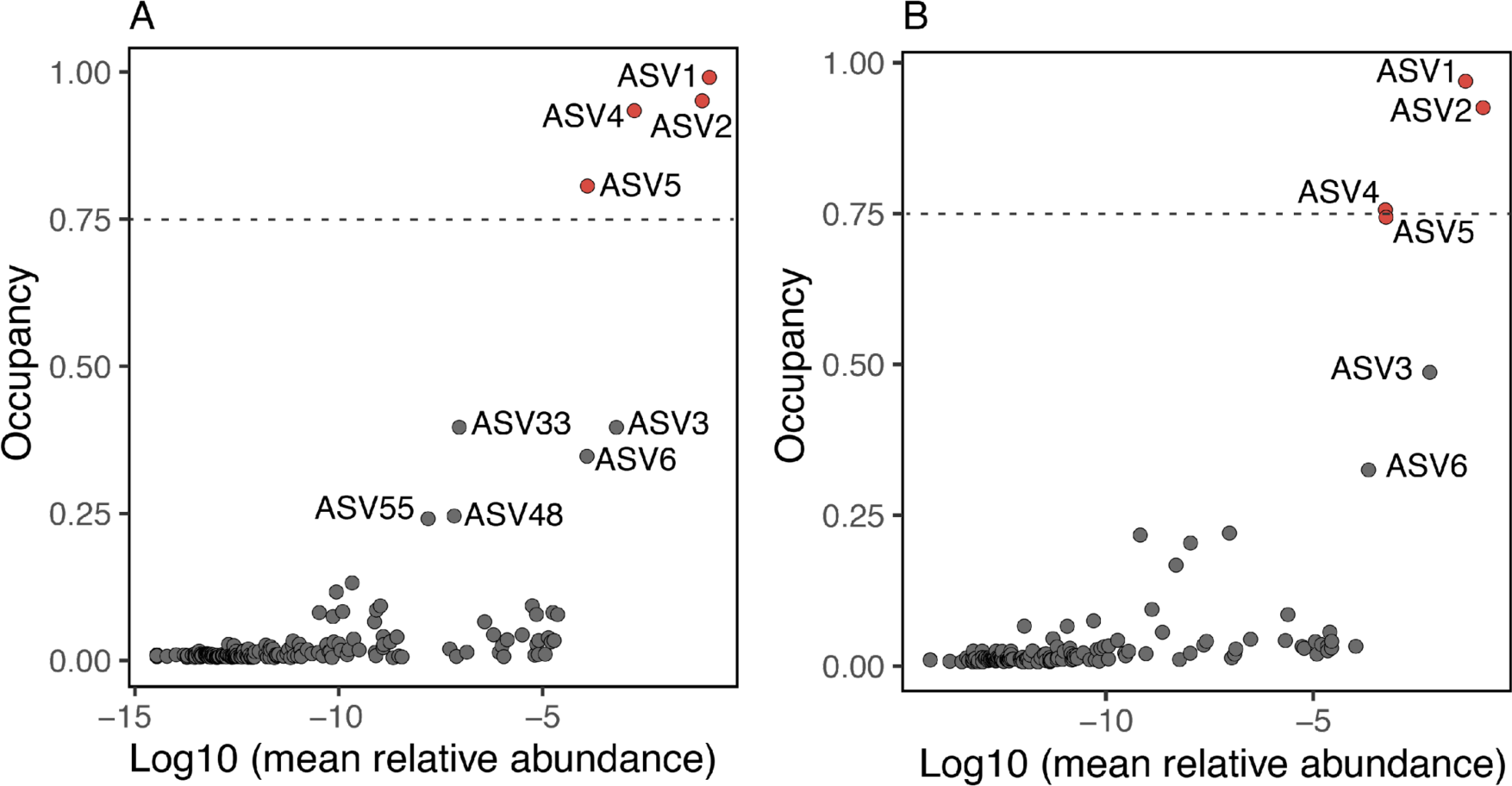
Abundance and Occupancy plot for ASVs for A) all five sites surveyed in 2021 (year 1) and B) Rowan and Union sites across two years (2021-22). ASVs highlighted in red have occupancy at or above 75%. ASVs identified with gray dots indicate occupancy greater than 25% and less than 75%.

## DISCUSSION

In this study, we characterized the wheat seed mycobiome in the context of local geographic structure, genotype, and spatial temporal variation. We used ITS1 metabarcoding to characterize the endophytic fungal community of wheat seeds and found that the seed mycobiome is largely resilient and stable in the context of local geography, genotype, and year. Some differences were observed for community species richness and composition by site. To obtain robust and fine scale definition of the wheat seed mycobiome, we examined the mycobiome of individual seeds. Given the gradients in latitude, longitude, soil types, and climates that the wheat seeds are exposed to in NC, we suggest that the stability of the wheat seed environment contributes to maintenance of a stable core group of seed associated taxa, while different environments allow for colonization of unique and rare taxa. Our findings represent a comprehensive assessment and investigation of the endophytic wheat seed mycobiome variation at the individual seed level that has only been previously reported in the context of disease (Bakker and McCormick 2019).

Across five sites in NC, we observed differences in fungal species richness for wheat seed samples from two cultivars collected in 2021. General correspondence between fungal community richness and geographical location has been previously established (Nicolaisen et al. 2014; Rojas et al. 2020; Sapkota et al. 2022). However, these studies assessed fungal communities of only two geographical sites and lacked robust replication at the site level. The differences in richness we observed can likely be attributed to the Washington field site’s coastal proximity and higher soil salinity (Table S1). Previous work by Mohamed and Martiny (2011) suggested that soil salinity drives fungal community composition of plant-associated endophytes when sampling across a salinity gradient on the US east coast.

When examining Poaceae fungal communities, *Alternaria alternata*, *Cladosporium* sp., and *Epicoccum nigrum* have been implicated in beneficial and mutualistic associations with host plants (Singh and Goodwin 2022; Wang et al. 2020; Kim et al. 2020; Fisher and Petrini 1992; DeMers 2022). Across species including wheat wild relatives and related grasses, genotypes, spatial and temporal gradients, these taxa were confirmed as core members of endophytic mycobiomes (Ofek-Lalzar et al. 2016; Özkurt et al. 2020; Comby et al. 2016; Duniere et al. 2017; Sapkota et al. 2022). Recently, *Alternaria* and *Cladosporium* sp. were reported to be dominant in the seed mycobiome of six forb species, and suggested to be vertically transmitted from pollen to seeds to cotyledons using a culture dependent approach (Hodgson et al. 2014). Our study provides additional evidence that these three fungal taxa constitute the core wheat seed mycobiome. We observed high occupancy rates for each taxon at or above ≥ 75% of samples, with each taxon present within every site and each cultivar and year sampled in addition to high relative abundance. Our findings provide evidence that these three taxa comprise the wheat seed core mycobiome.

Although *Alternaria*, *Epicoccum*, and *Cladosporium* play key roles by dominating the wheat seed mycobiome, several other taxa were notable for their slightly lower occupancy and relative abundance. Basidiomycota genera *Bullera*, *Filobasidium*, and *Papiliotrema* were among the moderately abundant taxa, which likely occur and grow as a yeast form within wheat seeds. Yeasts are considered a major component of above-ground wheat tissue communities, with *Bullera* and *Filobasidium* previously reported as inhabiting leaves and grains of cereals (Gouka et al. 2022; Nicolaisen et al. 2014; Solanki et al. 2021). We also detected the Ascomycete *Parastagonospora nodorum* which is well studied as the causal agent of Septoria Nodorum Blotch of wheat. Seed-borne transmission of this pathogen has been previously reported from wheat grown in the eastern United States (Cunfer 1978; Downie et al. 2021). Collectively, these moderately abundant taxa are often unique to a site or vary among samples within a single site and likely drive some of the variation that we observed at the community level within our study. While comparing fungal microbiome communities associated with two regionally adapted soft red winter wheat genotypes, we showed that genotypes did not contribute to differences in species richness or community composition. This finding supports previous studies that revealed little to no effect among wheat genotypes in assembly of fungal communities (Hassani et al. 2020). These observations are consistent with the current hypothesis that plant associated microbiomes should be structured by host plant phenotypes therefore, we expect that two cultivars with very similar traits in terms of class, heading, disease resistance, would exhibit comparable seed mycobiomes (Wagner 2021).

Seeds across many crop species have been characterized as simplistic and harboring low community diversity, with few taxa dominating as the most abundant and many rare taxa that may only be associated with a subset of samples. This common observation inspired the primary symbiont hypothesis that predicts few microbes dominate the seed endophytic tissue and play important functional roles in the developing seedling and/or adaptation of the plant host to the surrounding environment (Newcombe et al. 2018). To better understand the role that seed associated fungi play in shaping the seed microbial community and/or its function, we suggest an experimental approach that mimics the genomic biosurveillance of plant pathogens in plant tissues. This approach may involve whole genome sequencing of candidate fungal taxa, development of isolate-specific primers, and validation *in planta* to quantify and track the candidate fungal taxa in the seed and beyond.

Previous work on common beans suggested that single seeds were representative of plant-level variation with modest differences in fungal community composition among seeds from different individual plants (Bintarti et al. 2022). We aimed to capture mycobiome variation in individual plants within plots by sampling individual seeds. This sampling approach led to lower species richness estimates compared to previous studies examining the wheat seed mycobiome with pooled samples (Nicolaisen et al. 2014). When examining variation within our fungal community structure, the first two components of our PCoA accounted for 23% and 16% of variation respectively. Post-hoc analysis revealed that environmental and physiological factors such as latitude, longitude, seed moisture, yield, soil pH, and leaf nutrients (Ca, Fe, K, N, and Mn) accounted for 10-15% of variation in the seed mycobiome community composition. Therefore, a large component of fungal community variation is unaccounted for. However, this level of unexplained variance is typical of studies examining fungal mycobiomes of plant tissues (Lee and Hawkes 2021; Cregger et al. 2018; Leff et al. 2018). In our study, we observed high levels of variation in community composition within plots and within sites, this suggested that individual seeds mycobiomes remain largely stochastic. To reduce the stochastic effects of within plot and single seed variation, we hypothesize that sampling should occur at the individual plant level with corresponding plant level metadata collected. Among individual plant metadata, we consider it important to include seed nutrient analysis as the seed micro-environment may vary and impose selection on fungal taxa.

In conclusion, individual seed examination of the wheat seed mycobiome revealed a largely fixed fungal community in the context of local geography, genotype, and year. We found that the seed mycobiome is minorly shaped by site, which may be attributed to soil soluble salinity levels. Looking forward, it is critical for crop microbiome studies to take advantage of Official Variety Trials as they capture a spatial gradient of sites within a growing region and provide yield data and plot level metadata. These yield estimates, while not significant at explaining most of the variation seen in our studies, could provide a useful tool for identifying microbial community structure and taxa of interest in the future.

## ACKNOWLEDGMENTS

Thanks to David Marshall USDA-ARS Raleigh and Myron Fountain for providing seed for Hilliard. We are grateful to Gina Brown-Guedira, who provided seed for USG 3640. Thank you to our collaborators in the Collaborative Crop Resilience Program who provided valuable insight for this study. We thank Kevin Childs for their expertise and support for metabarcoding low-diversity plant tissues. We acknowledge the computing resources provided by North Carolina State University High Performance Computing Services Core Facility (RRID:SCR_022168). We thank Lisa Lowe for her guidance and support for using the computing services efficiently and Parker Ingraham for her invaluable help with skillful DNA extractions from isolates and seeds. This work was funded by NovoNordisk Fonden Grant #NNF19SA0059360.

---

**Table S1.** North Carolina State University Official Variety Trial site details, including latitude, longitude, rainfall in inches, soil type, plant date, harvest date, and soil salinity for 2021 and 2022.

| Site/Year | Latitude | Longitude | Rainfall<br>(in) | Soil Type | Plant<br>Date | Harvest<br>Date | Soil<br>salinity |
| --- | --- | --- | --- | --- | --- | --- | --- |
| **Lenoir** |  |  |  |  |  |  |  |
| 2021 | 35.37814 | -77.55983 | 42.48 | Rains sandy loam | 10/19/2020 | 6/14/2021 | 7.3 |
| **Robeson** |  |  |  |  |  |  |  |
| 2021 | 34.51122 | -79.30613 | 24.11 | Nahunta very fine sandy<br>loam | 11/23/2020 | 6/15/2021 | 6.1 |
| **Rowan** |  |  |  |  |  |  |  |
| 2021 | 35.69851 | -80.62480 | 31.59 | Lloyd clay loam | 10/22/2020 | 6/19/2021 | 7.3 |
| 2022 | 35.69649 | -80.62368 | 16.79 | Lloyd clay loam | 10/27/2021 | 6/7/2022 | 7.2 |
| **Union** |  |  |  |  |  |  |  |
| 2021 | 34.94934 | -80.42447 | 27.69 | Badin channery silt loam | 11/6/2020 | 6/19/2021 | 10 |
| 2022 | 34.85973 | -80.51312 | 22.42 | Cid channery silt loam | 11/3/2021 | 6/9/2022 | 10.5 |
| **Washington** |  |  |  |  |  |  |  |
| 2021 | 35.84917 | -76.65528 | 38.81 | Portsmouth fine sandy loam | 11/11/2020 | 6/16/2021 | 13.3 |

**Figure S1.**
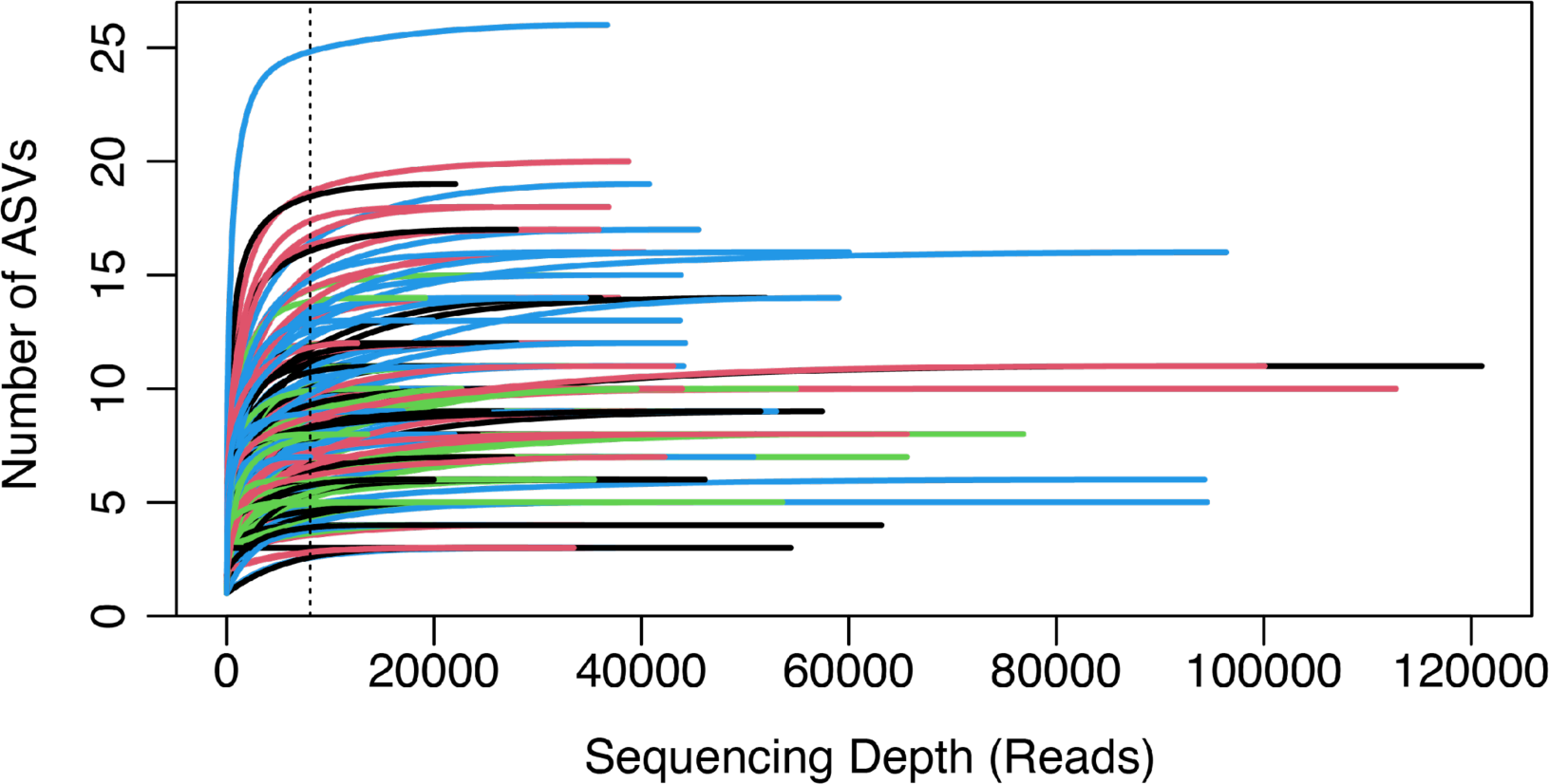
Sample rarefaction curve of 168 samples by read depth for metabarcoding. Vertical line indicates a minimum read depth of 8,076 reads.

**Figure S2.**
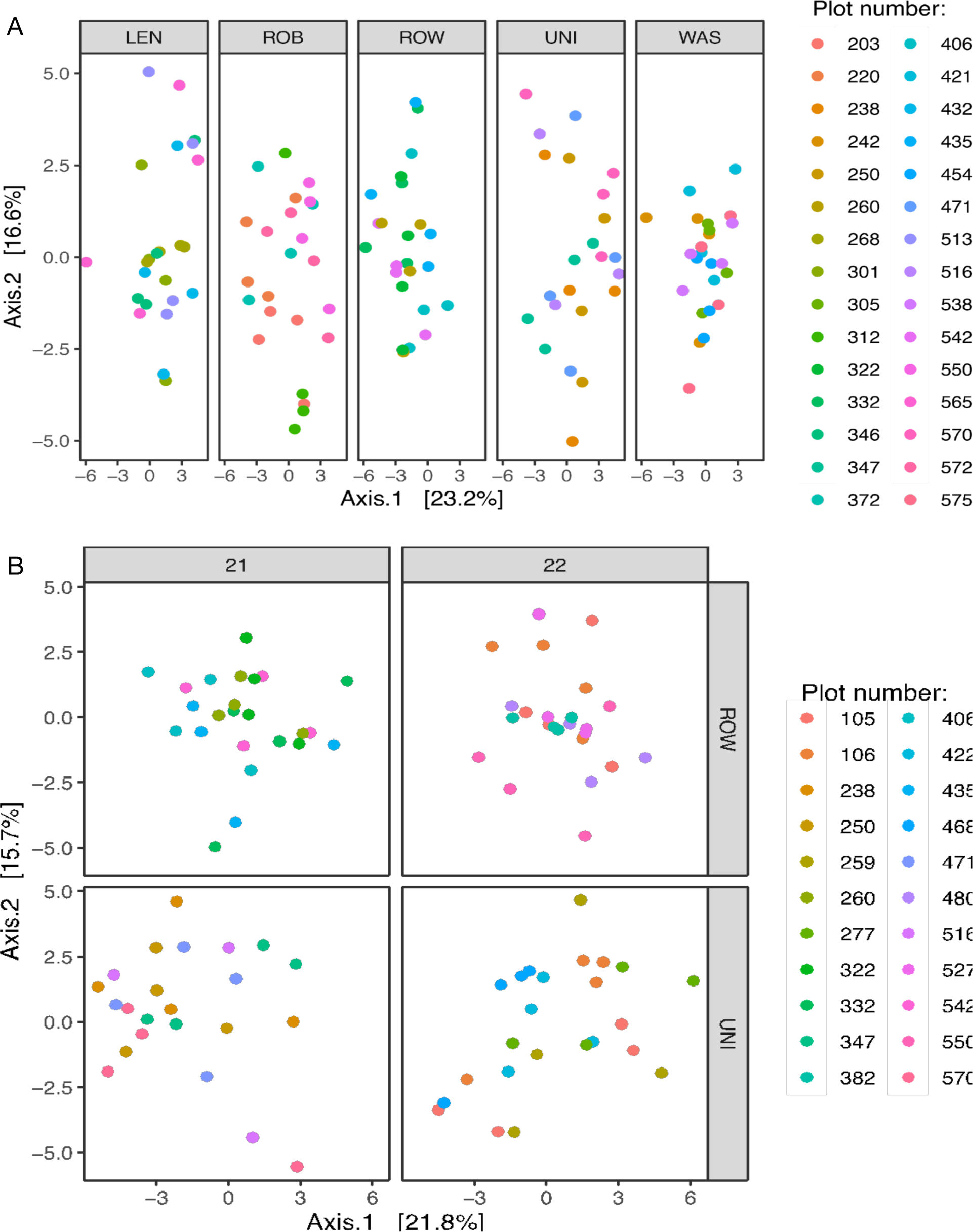
Principal Co-ordinate Analysis (PCoA) plot based on Euclidean Distances to examine structure of the wheat seed mycobiome. A) Seed mycobiome structure by site examining individual plot level variation for 2021 (year 1). B) Seed mycobiome structure for Rowan and Union experimental field sites examining individual plot level variation over two years from 2021-22.

## LITERATURE CITED

Abarenkov, K., Henrik Nilsson, R., Larsson, K.-H., Alexander, I. J., Eberhardt, U., Erland, S., et al. 2010. The UNITE database for molecular identification of fungi – recent updates and future perspectives. New Phytol. 186:281–285 Available at: http://onlinelibrary.wiley.com/doi/abs/10.1111/j.1469-8137.2009.03160.x

Above-Average-Multiple-Year_Commercial-2020_22.pdf. Available at: https://officialvarietytesting.ces.ncsu.edu/wp-content/uploads/2022/07/Above-Average-Multiple-Year_Commercial-2020_22.pdf?fwd=no.

Adam, E., Bernhart, M., Müller, H., Winkler, J., and Berg, G. 2018. The *Cucurbita pepo* seed microbiome: genotype-specific composition and implications for breeding. Plant Soil. 422:35–49 Available at: 10.1007/s11104-016-3113-9

Bakker, M. G., and McCormick, S. P. 2019. Microbial Correlates of Fusarium Load and Deoxynivalenol Content in Individual Wheat Kernels. Phytopathology. 109:993–1002 Available at: https://apsjournals.apsnet.org/doi/10.1094/PHYTO-08-18-0310-R

Barret, M., Briand, M., Bonneau, S., Préveaux, A., Valière, S., Bouchez, O., et al. 2015. Emergence Shapes the Structure of the Seed Microbiota. Appl. Environ. Microbiol. 81:1257–1266 Available at: https://journals.asm.org/doi/10.1128/AEM.03722-14

Benitez, M.-S., Osborne, S. L., and Lehman, R. M. 2017. Previous crop and rotation history effects on maize seedling health and associated rhizosphere microbiome. Sci. Rep. 7:15709 Available at: http://www.nature.com/articles/s41598-017-15955-9.

Bergelson, J., Mittelstrass, J., and Horton, M. W. 2019. Characterizing both bacteria and fungi improves understanding of the *Arabidopsis* root microbiome. Sci. Rep. 9:24 Available at: http://www.nature.com/articles/s41598-018-37208-z.

Bergna, A., Cernava, T., Rändler, M., Grosch, R., Zachow, C., and Berg, G. 2018. Tomato Seeds Preferably Transmit Plant Beneficial Endophytes. Phytobiomes J. 2:183–193 Available at: http://apsjournals.apsnet.org/doi/10.1094/PBIOMES-06-18-0029-R

Bintarti, A. F., Sulesky-Grieb, A., Stopnisek, N., and Shade, A. 2022. Endophytic Microbiome Variation Among Single Plant Seeds. Phytobiomes J. 6:45–55 Available at: https://apsjournals.apsnet.org/doi/full/10.1094/PBIOMES-04-21-0030-R

Borlaug, N. E. 1983. Contributions of Conventional Plant Breeding to Food Production. Science. 219:689–693 Available at: http://www.jstor.org/stable/1690517.

Callahan, B. J., McMurdie, P. J., Rosen, M. J., Han, A. W., Johnson, A. J. A., and Holmes, S. P. 2016. DADA2: High-resolution sample inference from Illumina amplicon data. Nat. Methods. 13:581–583.

Chaparro, J. M., Badri, D. V., and Vivanco, J. M. 2014. Rhizosphere microbiome assemblage is affected by plant development. ISME J. 8:790–803 Available at: http://www.nature.com/articles/ismej2013196.

Comby, M., Lacoste, S., Baillieul, F., Profizi, C., and Dupont, J. 2016. Spatial and Temporal Variation of Cultivable Communities of Co-occurring Endophytes and Pathogens in Wheat. Front. Microbiol. 7 Available at: https://www.frontiersin.org/articles/10.3389/fmicb.2016.00403/full

Cregger, M. A., Veach, A. M., Yang, Z. K., Crouch, M. J., Vilgalys, R., Tuskan, G. A., et al. 2018. The *Populus* holobiont: dissecting the effects of plant niches and genotype on the microbiome. Microbiome. 6:31.

Cunfer, B. M. 1978. The Incidence of *Septoria nodorum* in Wheat Seed. Phytopathology. 68:832 Available at: http://www.apsnet.org/publications/phytopathology/backissues/Documents/1978Abstracts/Phyto68_832.htm

DeMers, M. 2022. *Alternaria alternata* as endophyte and pathogen. Microbiol. Read. Engl. 168:001153.

Downie, R. C., Lin, M., Corsi, B., Ficke, A., Lillemo, M., Oliver, R. P., et al. 2021. *Septoria Nodorum* Blotch of Wheat: Disease Management and Resistance Breeding in the Face of Shifting Disease Dynamics and a Changing Environment. Phytopathology. 111:906–920 Available at: http://apsjournals.apsnet.org/doi/10.1094/PHYTO-07-20-0280-RVW

Duniere, L., Xu, S., Long, J., Elekwachi, C., Wang, Y., Turkington, K., et al. 2017. Bacterial and fungal core microbiomes associated with small grain silages during ensiling and aerobic spoilage. BMC Microbiol. 17:50 Available at: 10.1186/s12866-017-0947-0

Fisher, P. J., and Petrini, O. 1992. Fungal saprobes and pathogens as endophytes of rice (*Oryza sativa* L.). New Phytol. 120:137–143 Available at: http://onlinelibrary.wiley.com/doi/abs/10.1111/j.1469-8137.1992.tb01066.x.

Frøslev, T. G., Kjøller, R., Bruun, H. H., Ejrnæs, R., Brunbjerg, A. K., Pietroni, C., et al. 2017. Algorithm for post-clustering curation of DNA amplicon data yields reliable biodiversity estimates. Nat. Commun. 8:1188 Available at: http://www.nature.com/articles/s41467-017-01312-x

Gdanetz, K., and Trail, F. 2017. The Wheat Microbiome Under Four Management Strategies, and Potential for Endophytes in Disease Protection. Phytobiomes J. 1:158–168 Available at: https://apsjournals.apsnet.org/doi/10.1094/PBIOMES-05-17-0023-R

Gouka, L., Raaijmakers, J. M., and Cordovez, V. 2022. Ecology and functional potential of phyllosphere yeasts. Trends Plant Sci. 27:1109–1123 Available at: https://www.sciencedirect.com/science/article/pii/S1360138522001595

Griffey, C., Malla, S., Brooks, W., Seago, J., Christopher, A., Thomason, W., et al. 2020. Registration of ‘Hilliard’ wheat. J. Plant Regist. 14:406–417 Available at: http://onlinelibrary.wiley.com/doi/abs/10.1002/plr2.20073

Hassani, M. A., Özkurt, E., Franzenburg, S., and Stukenbrock, E. H. 2020. Ecological Assembly Processes of the Bacterial and Fungal Microbiota of Wild and Domesticated Wheat Species. Phytobiomes J. 4:217–224 Available at: http://apsjournals.apsnet.org/doi/10.1094/PBIOMES-01-20-0001-SC

Hertz, M., Jensen, I. R., Jensen, L. Ø., Thomsen, S. N., Winde, J., Dueholm, M. S., et al. 2016. The fungal community changes over time in developing wheat heads. Int. J. Food Microbiol. 222:30–39 Available at: https://www.sciencedirect.com/science/article/pii/S0168160516300198

Hodgson, S., Cates, C., Hodgson, J., Morley, N. J., Sutton, B. C., and Gange, A. C. 2014. Vertical transmission of fungal endophytes is widespread in forbs. Ecol. Evol. 4:1199–1208 Available at: https://www.ncbi.nlm.nih.gov/pmc/articles/PMC4020682/

Johnston-Monje, D., Lundberg, D. S., Lazarovits, G., Reis, V. M., and Raizada, M. N. 2016. Bacterial populations in juvenile maize rhizospheres originate from both seed and soil. Plant Soil. 405:337–355 Available at: 10.1007/s11104-016-2826-0.

Kim, H., Lee, K. K., Jeon, J., Harris, W. A., and Lee, Y.-H. 2020. Domestication of *Oryza* species eco-evolutionarily shapes bacterial and fungal communities in rice seed. Microbiome. 8:20 Available at: 10.1186/s40168-020-00805-0.

Klaedtke, S., Jacques, M.-A., Raggi, L., Préveaux, A., Bonneau, S., Negri, V., et al. 2016. Terroir is a key driver of seed-associated microbial assemblages. Environ. Microbiol. 18:1792–1804 Available at: https://onlinelibrary.wiley.com/doi/abs/10.1111/1462-2920.12977

Korbie, D. J., and Mattick, J. S. 2008. Touchdown PCR for increased specificity and sensitivity in PCR amplification. Nat. Protoc. 3:1452–1456 Available at: http://www.nature.com/articles/nprot.2008.133

Latz, M. A. C., Kerrn, M. H., Sørensen, H., Collinge, D. B., Jensen, B., Brown, J. K. M., et al. 2021. Succession of the fungal endophytic microbiome of wheat is dependent on tissue-specific interactions between host genotype and environment. Sci. Total Environ. 759:143804 Available at: https://www.sciencedirect.com/science/article/pii/S0048969720373356

Lee, M. R., and Hawkes, C. V. 2021. Plant and Soil Drivers of Whole-Plant Microbiomes: Variation in Switchgrass Fungi from Coastal to Mountain Sites. Phytobiomes J. 5:69–79 Available at: http://apsjournals.apsnet.org/doi/10.1094/PBIOMES-07-20-0056-FI.

Leff, J. W., Bardgett, R. D., Wilkinson, A., Jackson, B. G., Pritchard, W. J., De Long, J. R., et al. 2018. Predicting the structure of soil communities from plant community taxonomy, phylogeny, and traits. ISME J. 12:1794–1805 Available at: http://www.nature.com/articles/s41396-018-0089-x.

McMurdie, P. J., and Holmes, S. 2013. phyloseq: An R package for reproducible interactive analysis and graphics of microbiome census data. PLoS ONE. 8:e61217 Available at: http://dx.plos.org/10.1371/journal.pone.0061217.

Mergoum, M., Johnson, J., Buck, J., Buntin, G. D., Sutton, S., Lopez, B., et al. 2022. A new soft red winter wheat cultivar ‘GA 08535-15LE29’ adapted to Georgia and the U.S. southeast region. J. Plant Regist. 16:597–605 Available at: http://onlinelibrary.wiley.com/doi/abs/10.1002/plr2.20235

Mohamed, D. J., and Martiny, J. B. 2011. Patterns of fungal diversity and composition along a salinity gradient. ISME J. 5:379–388 Available at: http://www.nature.com/articles/ismej2010137

Nelson, E. B. 2018. The seed microbiome: Origins, interactions, and impacts. Plant Soil. 422:7–34 Available at: 10.1007/s11104-017-3289-7

Newcombe, G., Harding, A., Ridout, M., and Busby, P. E. 2018. A Hypothetical Bottleneck in the Plant Microbiome. Front. Microbiol. 9 Available at: https://www.frontiersin.org/articles/10.3389/fmicb.2018.01645

Nicolaisen, M., Justesen, A. F., Knorr, K., Wang, J., and Pinnschmidt, H. O. 2014. Fungal communities in wheat grain show significant co-existence patterns among species. Fungal Ecol. 11:145–153 Available at: https://www.sciencedirect.com/science/article/pii/S1754504814000907

Ofek-Lalzar, M., Gur, Y., Ben-Moshe, S., Sharon, O., Kosman, E., Mochli, E., et al. 2016. Diversity of fungal endophytes in recent and ancient wheat ancestors *Triticum dicoccoides* and *Aegilops sharonensis* ed. Linda Johnson. FEMS Microbiol. Ecol. 92:fiw152 Available at: https://academic.oup.com/femsec/article-lookup/doi/10.1093/femsec/fiw152

Oksanen, J., Simpson, G. L., Blanchet, F. G., Kindt, R., Legendre, P., Minchin, P. R., et al. 2022. vegan: Community Ecology Package. Available at: https://CRAN.R-project.org/package=vegan.

Özkurt, E., Hassani, M. A., Sesiz, U., Künzel, S., Dagan, T., Özkan, H., et al. 2020. Seed-Derived Microbial Colonization of Wild Emmer and Domesticated Bread Wheat (*Triticum dicoccoides* and *T. aestivum*) Seedlings Shows Pronounced Differences in Overall Diversity and Composition ed. Antonio Di Pietro. mBio. 11:e02637–20 Available at: https://journals.asm.org/doi/10.1128/mBio.02637-20.

Panke-Buisse, K., Poole, A. C., Goodrich, J. K., Ley, R. E., and Kao-Kniffin, J. 2015. Selection on soil microbiomes reveals reproducible impacts on plant function. ISME J. 9:980–989 Available at: http://www.nature.com/articles/ismej2014196.

Pebesma, E. 2018. Simple Features for R: Standardized Support for Spatial Vector Data. R J. 10:439–446 Available at: 10.32614/RJ-2018-009.

Peng, J. H., Sun, D., and Nevo, E. 2011. Domestication evolution, genetics and genomics in wheat. Mol. Breed. 28:281–301 Available at: 10.1007/s11032-011-9608-4.

R Core Team. 2018. R: A language and environment for statistical computing. R Foundation for Statistical Computing, Vienna, Austria. Available at: https://www.R-project.org.

Rojas, E. C., Sapkota, R., Jensen, B., Jørgensen, H. J. L., Henriksson, T., Jørgensen, L. N., et al. 2020. Fusarium Head Blight Modifies Fungal Endophytic Communities During Infection of Wheat Spikes. Microb. Ecol. 79:397–408 Available at: http://link.springer.com/10.1007/s00248-019-01426-3

Rybakova, D., Mancinelli, R., Wikström, M., Birch-Jensen, A.-S., Postma, J., Ehlers, R.-U., et al. 2017. The structure of the *Brassica napus* seed microbiome is cultivar-dependent and affects the interactions of symbionts and pathogens. Microbiome. 5:104 Available at: https://www.ncbi.nlm.nih.gov/pmc/articles/PMC5580328/

Sapkota, R., Jørgensen, L. N., Boeglin, L., and Nicolaisen, M. 2022. Fungal Communities of Spring Barley from Seedling Emergence to Harvest During a Severe *Puccinia hordei* Epidemic. Microb. Ecol. Available at: 10.1007/s00248-022-01985-y

Scibetta, S., Schena, L., Abdelfattah, A., Pangallo, S., and Cacciola, S. O. 2018. Selection and Experimental Evaluation of Universal Primers to Study the Fungal Microbiome of Higher Plants. Phytobiomes J. 2:225–236 Available at: http://apsjournals.apsnet.org/doi/10.1094/PBIOMES-02-18-0009-R

Shade, A., Jacques, M.-A., and Barret, M. 2017. Ecological patterns of seed microbiome diversity, transmission, and assembly. Curr. Opin. Microbiol. 37:15–22 Available at: https://www.sciencedirect.com/science/article/pii/S1369527416301576

Shade, A., and Stopnisek, N. 2019. Abundance-occupancy distributions to prioritize plant core microbiome membership. Curr. Opin. Microbiol. 49:50–58 Available at: https://www.sciencedirect.com/science/article/pii/S1369527419300426

Singh, R., and Goodwin, S. B. 2022. Exploring the Corn Microbiome: A Detailed Review on Current Knowledge, Techniques, and Future Directions. PhytoFrontiers^TM^. 2:158–175 Available at: http://apsjournals.apsnet.org/doi/10.1094/PHYTOFR-04-21-0026-RVW.

Smith, S. D. 2019. phylosmith: an R-package for reproducible and efficient microbiome analysis with phyloseq-objects. J. Open Source Softw. 4:1442 Available at: 10.21105/joss.01442.

Soil-and-Cultural-Practice-Table-2021.pdf. Available at: https://officialvarietytesting.ces.ncsu.edu/wp-content/uploads/2021/07/Soil-and-Cultural-Practice-Table-2021.pdf?fwd=no.

Soil-and-Cultural-Practice-Table-2022.pdf. Available at: https://officialvarietytesting.ces.ncsu.edu/wp-content/uploads/2022/07/Soil-and-Cultural-Practice-Table-2022.pdf?fwd=no.

Solanki, M. K., Abdelfattah, A., Sadhasivam, S., Zakin, V., Wisniewski, M., Droby, S., et al. 2021. Analysis of Stored Wheat Grain-Associated Microbiota Reveals Biocontrol Activity among Microorganisms against Mycotoxigenic Fungi. J. Fungi. 7:781.

Stopnisek, N., and Shade, A. 2021. Persistent microbiome members in the common bean rhizosphere: an integrated analysis of space, time, and plant genotype. ISME J. 15:2708–2722 Available at: http://www.nature.com/articles/s41396-021-00955-5.

Vannette, R. L. 2020. The Floral Microbiome: Plant, Pollinator, and Microbial Perspectives. Annu. Rev. Ecol. Evol. Syst. 51:363–386 Available at: 10.1146/annurev-ecolsys-011720-013401

Verma, S. K., Kharwar, R. N., Gond, S. K., Kingsley, K. L., and White, J. F. 2019. Exploring Endophytic Communities of Plants: Methods for Assessing Diversity, Effects on Host Development and Potential Biotechnological Applications. In Seed Endophytes: Biology and Biotechnology, eds. Satish Kumar Verma and Jr White James Francis. Cham: Springer International Publishing, p. 55–82. Available at: 10.1007/978-3-030-10504-4_4

Wagner, M. R. 2021. Prioritizing host phenotype to understand microbiome heritability in plants. New Phytol. 232:502–509 Available at: https://onlinelibrary.wiley.com/doi/abs/10.1111/nph.17622

Wang, M., Eyre, A. W., Thon, M. R., Oh, Y., and Dean, R. A. 2020. Dynamic Changes in the Microbiome of Rice During Shoot and Root Growth Derived From Seeds. Front. Microbiol. 11 Available at: https://www.frontiersin.org/articles/10.3389/fmicb.2020.559728.

Wickham, H. 2016. ggplot2: Elegant Graphics for Data Analysis. Springer-Verlag New York. Available at: https://ggplot2.tidyverse.org.

Wolfgang, A., Zachow, C., Müller, H., Grand, A., Temme, N., Tilcher, R., et al. 2020. Understanding the Impact of Cultivar, Seed Origin, and Substrate on Bacterial Diversity of the Sugar Beet Rhizosphere and Suppression of Soil-Borne Pathogens. Front. Plant Sci. 11:560869 Available at: https://www.ncbi.nlm.nih.gov/pmc/articles/PMC7554574/

